# Protection from lethal *Clostridioides difficile* infection via intraspecies competition for co-germinant

**DOI:** 10.1101/2021.02.26.433146

**Authors:** Jhansi L. Leslie, Matthew L. Jenior, Kimberly C. Vendrov, Alexandra K. Standke, Madeline R. Barron, Tricia J. O’Brien, Lavinia Unverdorben, Pariyamon Thaprawat, Ingrid L. Bergin, Patrick D. Schloss, Vincent B. Young

**Author notes:** Jhansi L. Leslie, Department of Medicine, Division of Infectious Diseases and International Health, University of Virginia, Charlottesville, Virginia, USA; Matthew L. Jenior, Department of Biomedical Engineering, University of Virginia, Charlottesville, Virginia, USA; Tricia O’Brien, College of Osteopathic Medicine, Michigan State University, East Lansing, Michigan, USA.

## Abstract

*Clostridioides difficile,* a Gram-positive, spore-forming bacterium, is the primary cause of infectious nosocomial diarrhea. Antibiotics are a major risk factor for *C. difficile* infection (CDI) as they disrupt the gut microbial community, enabling increased germination of spores and growth of vegetative *C. difficile*. To date the only single-species bacterial preparation that has demonstrated efficacy in reducing recurrent CDI in humans is non-toxigenic *C. difficile.* Using multiple infection models we determined that pre-colonization with a less virulent strain is sufficient to protect from challenge with a lethal strain of *C. difficile,* surprisingly even in the absence of adaptive immunity. Additionally, we showed that protection is dependent on high levels of colonization by the less virulent strain and that it is mediated by exclusion of the invading strain. Our results suggest that reduction of amino acids, specifically glycine following colonization by the first strain of *C. difficile* is sufficient to decrease germination of the second strain thereby limiting colonization by the lethal strain.

**Importance:** Antibiotic-associated colitis is often caused by infection with the bacterium *Clostridioides difficile.* In this study we found that reduction of the amino acid glycine by pre-colonization with a less virulent strain of *C. difficile* is sufficient to decrease germination of a second strain. This finding demonstrates that the axis of competition for nutrients can include multiple life stages. This work is important, as it is the first to identify a possible mechanism through which pre-colonization with *C. difficile*, a current clinical therapy, provides protection from reinfection. Furthermore, our work suggests that targeting nutrients utilized by all life stages could be an improved strategy for bacterial therapeutics that aim to restore colonization resistance in the gut.

## Introduction

*Clostridioides difficile,* a Gram-positive, spore-forming bacterium, is the primary cause of infectious nosocomial diarrhea (1). Susceptibility to *C. difficile* infection (CDI) results from perturbations of the gut microbial community, enabling increased germination of spores and growth of vegetative *C. difficile* (2, 3). Following colonization, vegetative *C. difficile* produces the toxins TcdA and TcdB, which are glycosyltransferases that inactivate cellular GTPases (4). Inactivation of these key cellular proteins results in damage of the colonic epithelium and inflammation manifesting as colitis, diarrhea and in severe cases, toxic megacolon or even death. Currently the principal treatment for acute CDI is the antibiotic vancomycin (5). While this treatment limits vegetative *C. difficile*, this therapy further disrupts the gut microbiota delaying community recovery, potentially leading to recurrent disease (6).

Due to the high rate of recurrent infection associated with existing treatment for CDI, alternative approaches that spare or even restore the gut microbiota have been a focus of recent work. As is the case with many toxin-mediated diseases, early studies noted that generation of a humoral immune response to the toxins could be sufficient to protect against disease (7). A recent clinical trial demonstrated that patients receiving monoclonal antibodies targeting the toxins were fifty percent less likely to experience recurrent disease (8, 9). Unlike antibiotics, antibody therapy prevents illness while likely sparing the microbial community. Recently, a non-toxigenic strain of *C. difficile* was demonstrated to successfully reduce the rate of recurrent CDI by approximately fifty percent (10, 11). The prevailing hypothesis is that the protection provided by the non-toxigenic strain is mediated by competitive exclusion, limiting the ability of toxigenic *C. difficile* to colonize the gut (12). However, this has never been conclusively demonstrated. Using a murine model of CDI, we sought to address this question.

We report here that, in the absence of adaptive immunity, pre-colonization with one strain of *C. difficile* is sufficient to provide protection from lethal infection with another. Using gnotobiotic mice we show that protection is mediated by limiting colonization of the highly virulent strain. Furthermore, we provide evidence that exclusion is not predicated on nutrient-based limitation of the vegetative form of the invading strain, but rather on depletion of the co-germinant glycine. This work is important, as it is the first study to identify a possible mechanism through which pre-colonization with *C difficile*, a current clinical therapy, provides protection from recurrent CDI. Furthermore, limitation of germination due to decreased levels of glycine in the gut is a novel paradigm for colonization resistance.

## Results

### Development of a murine model of persistent *C. difficile* colonization

The only single species bacterial preparation that has demonstrated efficacy in reducing recurrent CDI in humans is non-toxigenic *C. difficile* (10). However the mechanisms by which one strain of *C. difficile* prevents colonization of another are currently unknown. To begin to address this question we developed a model of persistent *C. difficile* colonization. Mice were made susceptible to colonization via administration of the antibiotic cefoperazone. Following two days off of the antibiotic, mice were either mock challenged or challenged with *C. difficile* strain (str.) 630 spores. Compared to mock-challenged animals, infected animals displayed significant weight loss between days four to six post-challenge (figure 1A, *P* < 0.05). After 10 days, infected mice remained stably colonized at approximately 10^7^ CFU/g feces despite a significant reduction in fecal levels of *C. difficile* relative to levels on day one post-infection (figure 1 B, *P* < 0.01). Coincident with the decrease in colonization, infected animals recovered weight until they were indistinguishable from the mock-infected controls (figure 1A). Additionally, the diversity of the gut microbiota increased within the first 10 days of infection corresponding to the recovery of weight and decrease in pathogen colonization (figure 1C box plots, right axis). Throughout the experiment, toxin was detected in the feces of most of the infected mice. However, toxin activity started to decrease by day 14 and was significantly decreased by day 33 post challenge relative to early in the infection (figure 1D). Together these data demonstrate that *C. difficile* str. 630 can persistently colonize wild type mice as a minority member of the gut microbiota (figure 1C, left axis).

**Figure 1:**
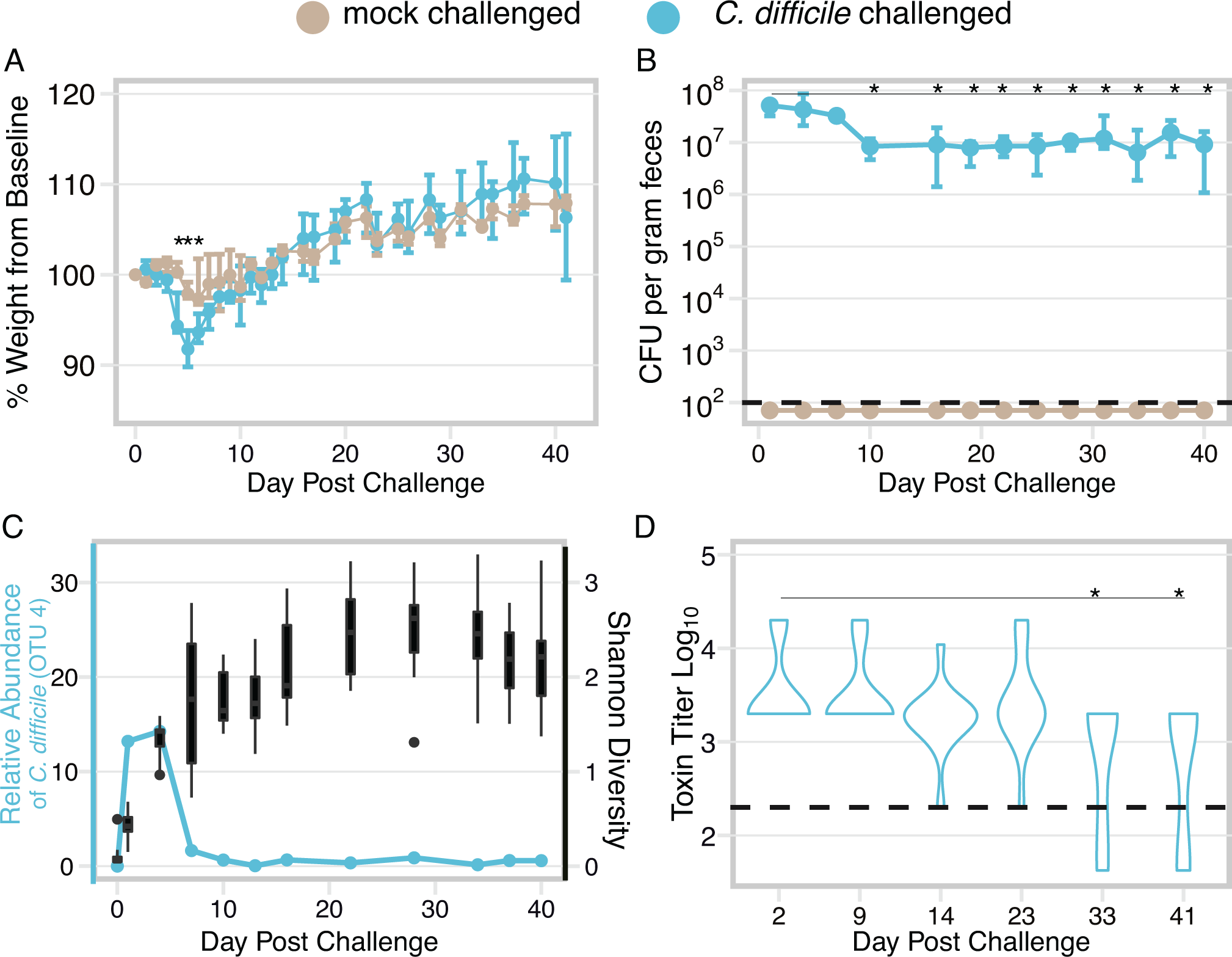
Murine model of persistent *C. difficile* colonization. **A.** Change in weight relative to day of infection in infected and mock challenged mice. Points represent median weight; bars are the upper and lower quartiles. Infected mice are colored blue (n=14) while data from mock-infected animals are shown in tan (n=8). Following correction for multiple comparisons, weight loss in infected mice was only significantly different than mock-challenged mice on days 4, 5, and 6 post-infection, *P* < 0.05. **B.** *C. difficile* colonization over time as determined by quantitative culture. Colonization significantly decreased and remained significantly lower by day 10 post-infection relative to day 1 post-infection 40 (n=14), *P* < 0.01. The dashed line represents the limit of detection of 100 CFU/g feces. **C.** Relative abundance of OTU 4 (*C. difficile*) over the course of the experiment (blue line) is plotted on the left axis while Shannon diversity of the infected mice over the course of the experiment is plotted on the right axis (black box plots). **D.** Fecal toxin activity remains detectable throughout the experiment. Toxin titers on day 33 and day 40 are significantly different from day one post-infection levels (n=14), *P* < 0.05. Statistical significance for all data was calculated using a Wilcoxon test with Benjamin-Hochberg correction. The dashed line represents the limit of detection (LOD) for each assay, for visual clarity samples that were below the limit of detection were plotted below the line. However, for statistical analysis, the value of LOD/√2 was substituted for undetected values.

### Pre-colonization with *C. difficile* protects mice from challenge with a highly virulent strain even in the absence of adaptive immune responses

To determine if a resident strain of C*. difficile* protects mice from challenge with a second strain in the context of perturbation to the gut microbiota, we administered a second antibiotic, clindamycin, to both the colonized and un-colonized mice. Previous work from our group demonstrated that a single dose of clindamycin rendered mice susceptible to colonization following six-weeks of recovery from cefoperazone treatment (13). Clindamycin did not result in weight loss in either group of mice (Supplemental figure 1A). However, levels of str. 630 in the colonized mice significantly increased following administration of clindamycin likely because it is resistant to the antibiotic (Supplemental figure 1B, *P*< 0.001) (14). The day after mice were given clindamycin, they were challenged with 10^5^ CFU of str. VPI 10463 spores. Strain VPI 10463 is lethal in this model of infection (13). To control for possible variations in the microbiota across cages, mice from str. 630-infected and mock-infected (naïve) cages were split into two groups (figure 2A). Approximately half the mice in a cage were challenged with str. VPI 10463 while the remaining mice were placed in a new cage and received mock infection. Naïve mice that had neither been infected with str. 630 nor str. VPI 10463 maintained stable weight, however naïve mice that were challenged with str. VPI 10463 lost a significant amount of weight and had to be euthanized (figure 2B, *P* < 0.01). Mice that were persistently colonized with str. 630 did not lose weight despite being challenged with the lethal strain. In addition, str. 630 pre-colonized mice had significantly lower toxin titers than the naïve mice challenged with str. VPI 10463 (figure 2C, *P* < 0.05). This finding was confirmed by histopathology, as the summary scores of the histopathological damage in the colon was significantly less in the str. 630 colonized mice challenged with str. VPI 10463 compared to the naïve mice challenged with str. VPI 10463 (figure 2D *P* < 0.01).

**Figure 2:**
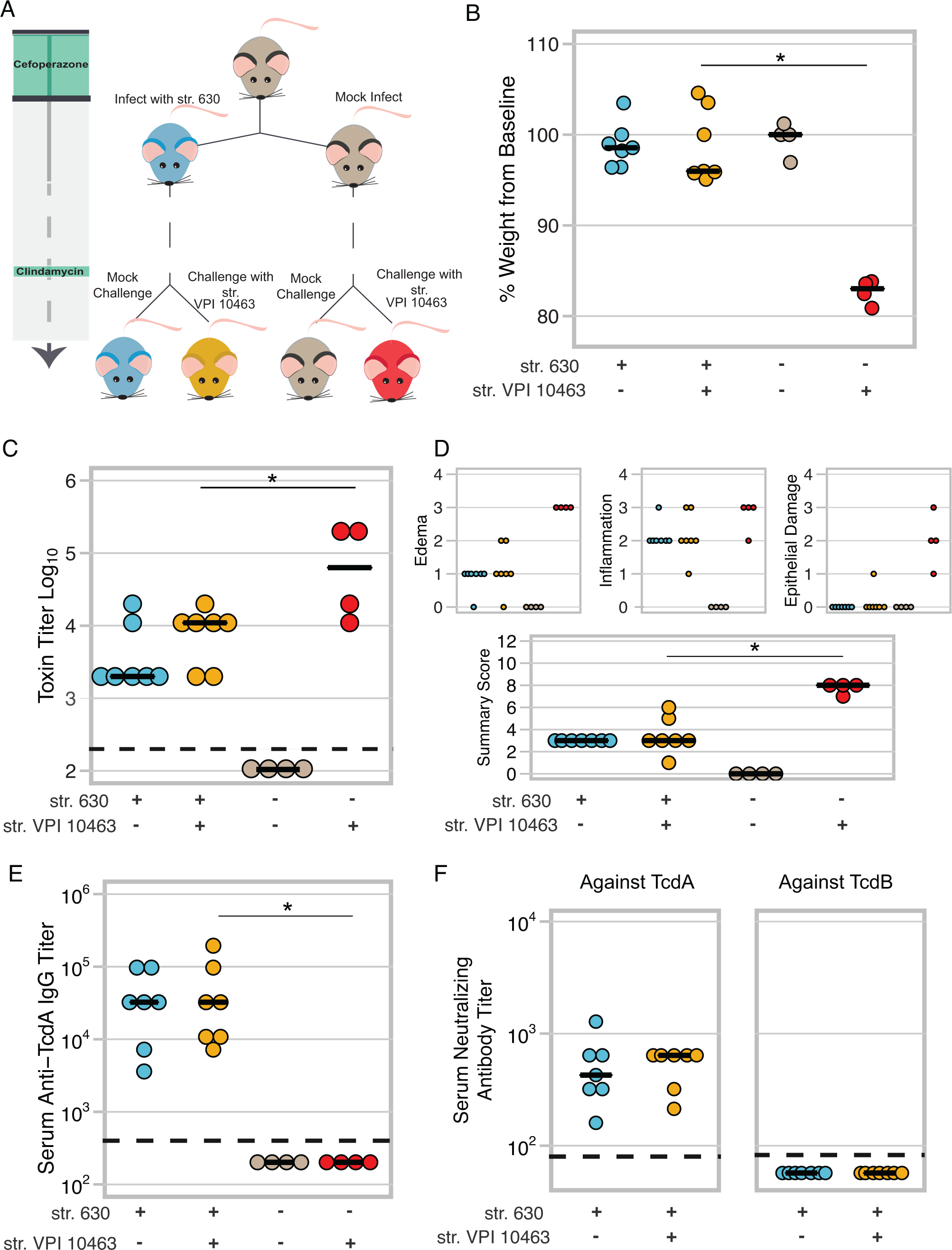
Mice pre-colonized with *C. difficile* str. 630 are protected from challenge with a lethal strain. **A.** Schematic of experimental conditions. Colors corresponding to treatment groups are carried throughout the figure. **B.** Change in weight at time of necropsy relative to weight on day of challenge, day 44 of the model. Mice colonized with *C. difficile* str. 630 and challenged with the lethal strain (VPI 10463) (n=4) are protected from weight loss whereas mice that had no exposure to *C. difficile* str. 630 experienced significant weight loss (n=4), *P* < 0.01. **C.** Toxin titer from intestinal content from mice in panel A as measured by Vero cell rounding assay. Mice colonized with *C. difficile* str. 630 and then challenged with *C. difficile* str. VPI 10463 have a lower toxin titer relative to naïve mice challenged with str. VPI 10463, *P* < 0.05. **D.** Histopathology from colons of the mice in panel A. Small panels depict scores for each component of the summary score. Summary score of str. VPI 10463 challenged str. 630 colonized vs. str. VPI 10463 challenged naive mice *P* < 0.01. **E.** Titer of serum IgG against TcdA at conclusion of experiment as measured by ELISA. Limit of detection was a titer of 400, *P* < 0.01. **F.** Neutralizing titer of serum against TcdA or TcdB. For all data statistical significance between the *C. difficile* str. VPI 10463 challenged str. 630-colonized and str. VPI 10463 challenged naive mice was determined by Wilcoxon test. The dashed line represents the limit of detection for each assay, for visual clarity samples that were below the limit of detection were plotted below the line.

Since both strains express nearly identical forms of TcdA and TcdB and we observed decreased signs of disease in the primary infection after a week of first being challenged, we questioned if protection might be due to the development of a humoral immune response to the toxins (7, 15). We found that mice previously infected with str. 630 developed a high anti-TcdA titer with a median titer of 1:32,400 with a portion of these antibodies being neutralizing, capable of preventing TcdA mediated cell rounding (figure 2E str. 630 colonized vs. naïve mice *P* < 0.01 & 2F).

Since protection was correlated with both pre-colonization and the development of an adaptive immune response to the toxins, we next tested if adaptive immunity was the sole factor preventing disease in our model by utilizing mice defective in recombination-activating gene 1 (RAG), a gene that is critical in the development of B and T cells. RAG1^-/-^ lack the adaptive arm of their immune system (16). Following 40 days of colonization, mice were given an intraperitoneal injection of clindamycin and then challenged with spores of the lethal str. VPI 10463. Surprisingly, both the RAG^-/-^ and WT mice pre-colonized with str. 630 were protected following the challenge, while the naïve mice of both genotypes succumbed to the infection (figure 3A and B, naïve vs. colonized *P* < 0.05). Scoring of the colonic pathology demonstrated that both the RAG1^-/-^ and WT mice pre-colonized with str. 630 had less pathology relative to the naïve mice (figure 3C, naïve vs. colonized *P* < 0.05). These data demonstrate that in the absence of adaptive immunity pre-colonization with *C. difficile* is sufficient to protect from lethal infection with spores of another strain. Furthermore, protection from severe disease in mice pre-colonized with str. 630 is not mediated by adaptive immunity.

**Figure 3:**
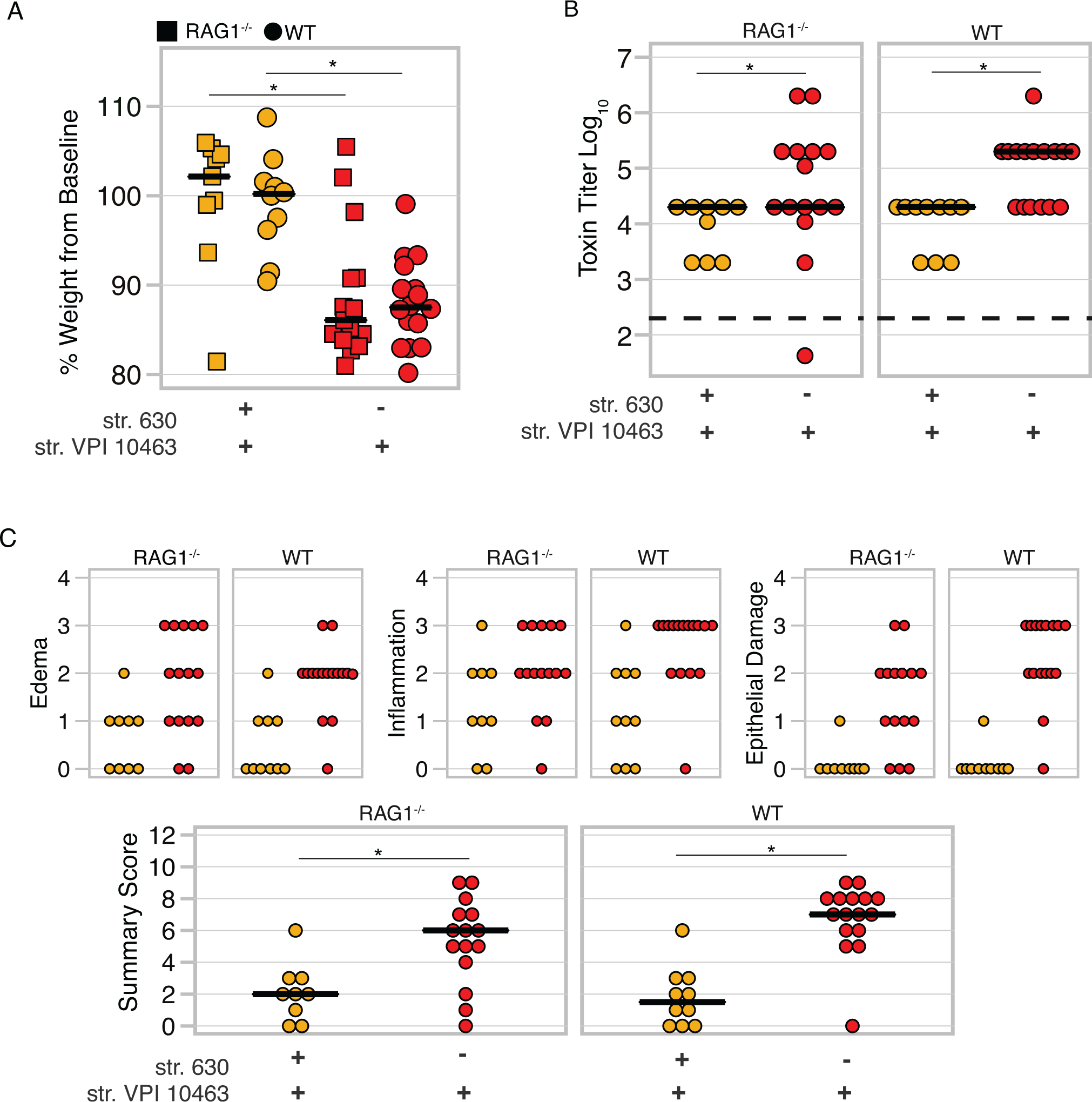
RAG1^-/-^ mice pre-colonized with *C. difficile* are protected from challenge with a lethal strain of *C. difficile*. **A.** Change in weight at time of necropsy relative to weight on day of challenge, day 44 of the model. Both WT and RAG1^-/-^ mice colonized with str. 630 and then challenged with the highly virulent str. VPI 10463 are protected from weight loss whereas mice that had no exposure to str.630 experienced significant weight loss, str. VPI 10463 challenged str. 630 colonized RAG1^-/-^ vs. VPI challenged naive RAG1^-/-^ p <0.05, str. VPI 10463 challenged str. 630 colonized WT vs. str. VPI 10463 challenged naive WT *P* < 0.01, no statistical difference was detected in comparisons of the same treatment between the two genotypes. **B.** Toxin titer from intestinal content of mice in figure A as measured by Vero cell cytotoxicity assay. Both WT and RAG1^-/-^ mice colonized with str. 630 and then challenged with str. VPI 10463 have a lower toxin titer relative to naïve mice challenged with str. VPI 10463. Str. VPI 10463 challenged and str. 630 colonized RAG1^-/-^ vs. str. VPI 10463 challenged naive RAG1^-/-^ *P* < 0.05. Str. VPI 10463 challenged and str. 630 colonized WT vs. str. VPI 10463 challenged naive WT *P* < 0.001. Statistical significance was calculated using a Wilcoxon test. Limit of detection was 2.3, however for visual clarity samples with an undetected toxin titer were plotted below the limit of detection. **C.** Histopathology scoring of the colon damage of mice from panel A. Smaller panels depict scores for each component of the summary score. Summary scores of str. VPI 10463 challenged str. 630 colonized RAG1^-/-^ vs. str. VPI 10463 challenged naive RAG1^-/-^ *P* < 0.05, str. VPI 10463 challenged str. 630 colonized WT vs. str. VPI 10463 challenged naive WT *P* < 0.001. For all panels, statistical significance was calculated using a Wilcoxon test with Benjamini-Hochberg corrections when appropriate. Data are from two independently run experiments with multiple cages per each treatment group.

### Protection afforded by low virulence *C. difficile* strain develops rapidly and depends on limiting colonization of the lethal strain

Having excluded the contribution of the adaptive immune response in our model, we next tested if protection required treatment with live *C. difficile*. Activation of innate immune pathways with microbe-associated molecular patterns, such Toll-like receptor 5 with flagellin, protects against acute CDI (17, 18). Additionally, in other colitis models, both viable and heat-killed probiotic strains ameliorate disease via stimulation of host innate immune pathways (19, 20). Thus, we tested if defense against severe CDI could be conferred by pre-treatment of mice with a high dose of heat-killed vegetative str. 630. As we had not observed a role for adaptive immunity we shortened the model from 42 days of pre-infection with str. 630 to one day. However, since we were additionally testing heat-killed vs. live colonization we included RAG1^-/-^ mice in this experiment to confirm findings from our persistent colonization model. Cefoperazone-treated mice were challenged with str. 630 spores, given the equivalent of 10^9^ CFU of autoclaved str. 630, or mock infected. 24 hours (hrs.) later all mice were challenged with str. VPI 10463 spores (Supplemental figure 2A). We observed no protective effect of heat-killed str. 630, indicating that protection requires colonization with *C. difficile* str. 630 not merely exposure to antigen (figure 4A, *P* < 0.05 for all comparisons where significance is indicated). In this short infection model, we repeated our finding that adaptive immunity is not necessary for protection as both RAG1^-/-^ and WT mice colonized with str. 630 were protected from weight-loss after just 24 hrs. of colonization. Mice given viable *C. difficil*e str. 630 were highly colonized with the strain, while str. 630 was not detected in mice that received the heat-killed str. 630 or mock infection (figure 4B). Total levels of *C. difficile* were not significantly different between the groups (figure 4C, *P* > 0.7). These data suggested that pre-colonization with str. 630 protects by limiting colonization of str. VPI 10463.

**Figure 4:**
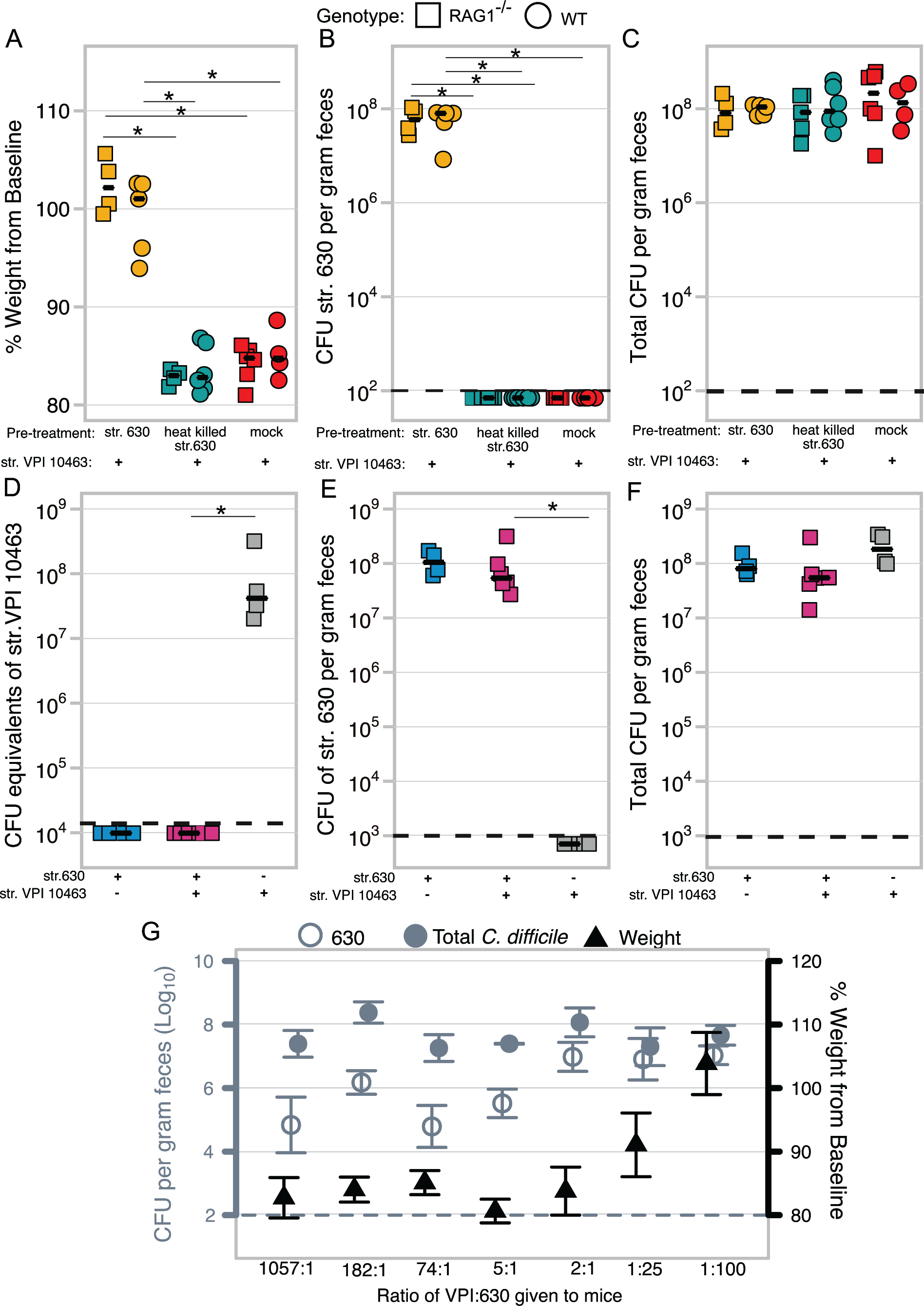
*C. difficile* str. 630 protects by limiting colonization of the lethal strain. **A.** Change in weight at time of necropsy (day 3) relative to weight on day of challenge in mice pre-treated with viable str.630, heat-killed str. 630 or water (mock). All mice were infected with str. VPI 10463 one day following pre-treatment. Both RAG1^-/-^ or WT mice given viable str. 630 did not lose weight following challenge with str. VPI 10463 when compared to mice who received heat-killed str. 630 or mock, *P* < 0.05 for all comparisons shown. **B**. Levels of str. 630 in mice at conclusion of experiment on day three (str. 630 is erythromycin resistant while str. VPI 10463 is sensitive to the antibiotic). Colonization by str. 630 was significantly different in mice given viable str. 630 compared to mice given heat-killed str. 630 or mock. RAG1^-/-^ mice given str. 630 vs. heat-killed str. 630, *P* < 0.05 or vs. mock, *P* < 0.05. WT mice given str. 630 vs. heat-killed str. 630, *P* < 0.05 or vs. mock, *P* < 0.05. There was no significant difference between colonization in the mock vs. heat-killed str. 630 treatments for either genotype. LOD is 100 CFU/g feces; undetected samples were plotted below the LOD for visual clarity. **C.** Total levels of *C. difficile* colonization at conclusion of experiment. There was no significant difference in total *C. difficile* colonization between any of the groups, *P* > 0.7. **D.** CFU equivalents of str. VPI 10463 in gnotobiotic mice as determined by qPCR on day three. Mice pre-colonized with *C. difficile* str. 630 have undetectable levels of str. VPI 10463 using this assay, LOD is 1.39 x10^4^ CFU, levels of str. VPI 10463 in str. 630 pre-colonized mice vs. str. VPI 10463 only mice *P* < 0.01. **E**. CFU/gram of feces of *C. difficile* strain str. 630 in gnotobiotic across groups as determined by selective quantitative culture. Mice only challenged with str. VPI 10463 were not colonized with str. 630. LOD is 1000 CFU, *P* < 0.05. **F.** Total CFU/gram of feces of *C. difficile* in gnotobiotic as determined by quantitative culture on day three. **G.** Mice challenged simultaneously with both strains can be colonized by both strains. Left axis represents Log_10_ CFU of total *C. difficile* (closed circle) or str. 630 two days post challenge (open circle). Right axis depicts percent of baseline weight two days post challenge (triangle). Each different inoculum ratio was given to one cage of five mice. Points represent the median value for each treatment while the bars represent the upper and lower quartiles. For all panels in the figure, squares represent RAG1^-/-^ mice while circles represent wild-type (WT) mice. For the data included in each figure, statistical significance was calculated using a Wilcoxon test and corrected with a Benjamini-Hochberg correction. The dashed line represents the limit of detection for each assay, for visual clarity samples that were below the limit of detection were plotted below the line.

Since treatment with viable *C. difficile* was required for protection, we sought to establish if this was mediated directly by str. 630, rather than indirectly through changes in the microbiota. To test this, we utilized gnotobiotic mice. Using our selective plating scheme, it is only possible to differentiate str. 630 from total *C. difficile* as we were unable to identify an antibiotic that only str. VPI 10463 was resistant to. To overcome this limitation, we developed a quantitative PCR assay using primers that amplify a target in str. VPI 10463 that is absent in str. 630 (Supplemental figures 3A and B). While these primers are not specific to solely str. VPI 10463 when used in the context of the rest of the gut microbiota, they could be used in gnotobiotic mice. RAG1^-/-^ germ-free mice were either infected with str. 630 spores or left germ-free; the following day, the germ-free mice in addition to one of the groups of str.630-monoassociated mice were challenged with str. VPI 10463 spores (Supplemental figure 2B). Using this model, we were able to determine that pre-colonization with str. 630 is sufficient to prevent colonization by spores of the lethal strain, as we were unable to detect str. VPI 10463 genomic DNA in the mice that were co-challenged despite high overall levels of *C. difficile* measured by quantitative culture (figures 4D, E, F).

Reports of patients infected with multiple strains of *C. difficile* suggest that despite our results, infection with multiple strains can occur (21, 22). When we infected mice with different ratios of spores from each of the two strains simultaneously, we found that both strains were capable of colonizing despite starting with over a 1000x less str. 630 (Supplemental figure 2C). While both strains were detectable when co-inoculated, protection, defined as a lack of weight-loss and low clinical scores, was not observed unless str. 630 was the dominant strain (figure 4G, Supplemental figures 4A-D). Interestingly, when infecting with different ratios of the two strains, the total burden would not pass a threshold of 10^9^, suggesting a population carrying capacity for *C. difficile* in the mouse gut. Together these data demonstrate that protection from lethal disease requires colonization with high levels of str. 630 to prevent establishment of the second strain.

### Limitation of the lethal strain is mediated by decreased availability of a co-germinant, glycine

Others have reported that pre-colonization with one strain of *C. difficile* provides protection from challenge with a more virulent strain (23–26). The prevailing hypothesis is that consumption of nutrients by the first strain limits the ability of the invading strain to grow (12). We tested this hypothesis in an *ex vivo* assay using sterile medium prepared from the cecal contents of a susceptible mice. Using this approach, we found that when vegetative *C. difficile* was inoculated into susceptible mouse cecal medium, both strains displayed significant growth after 24 hrs. (Supplemental figure 5A, *P* < 0.001). To test if one day of colonization was sufficient to reduce the nutrients required for growth, we added vegetative str. VPI 10463 to filter sterilized spent culture from the experiments in Supplemental figure 3A. Spent cecal medium from 24 hrs. cultures of both str. 630 and str. VPI 10463 supported another round of significant growth (figure 5A). To test if nutrient utilization by str. 630 was different *in vivo,* we additionally assessed growth of str. VPI 10463 in cecal medium made from mice infected with str. 630 for 24 hrs. This medium also supported robust growth, demonstrating that in both batch culture and *in vivo*, 24 hrs. of colonization is not sufficient to reduce the nutrients required for growth of a vegetative invading strain (Supplemental figure 5B). Additionally, as we were able to culture vegetative str. VPI 10463 in the spent culture of str. 630, we excluded the possibility of inhibition due to secreted products like bacteriocins or phage in this model. This was confirmed by agar overlay assays (data not shown).

**Figure 5:**
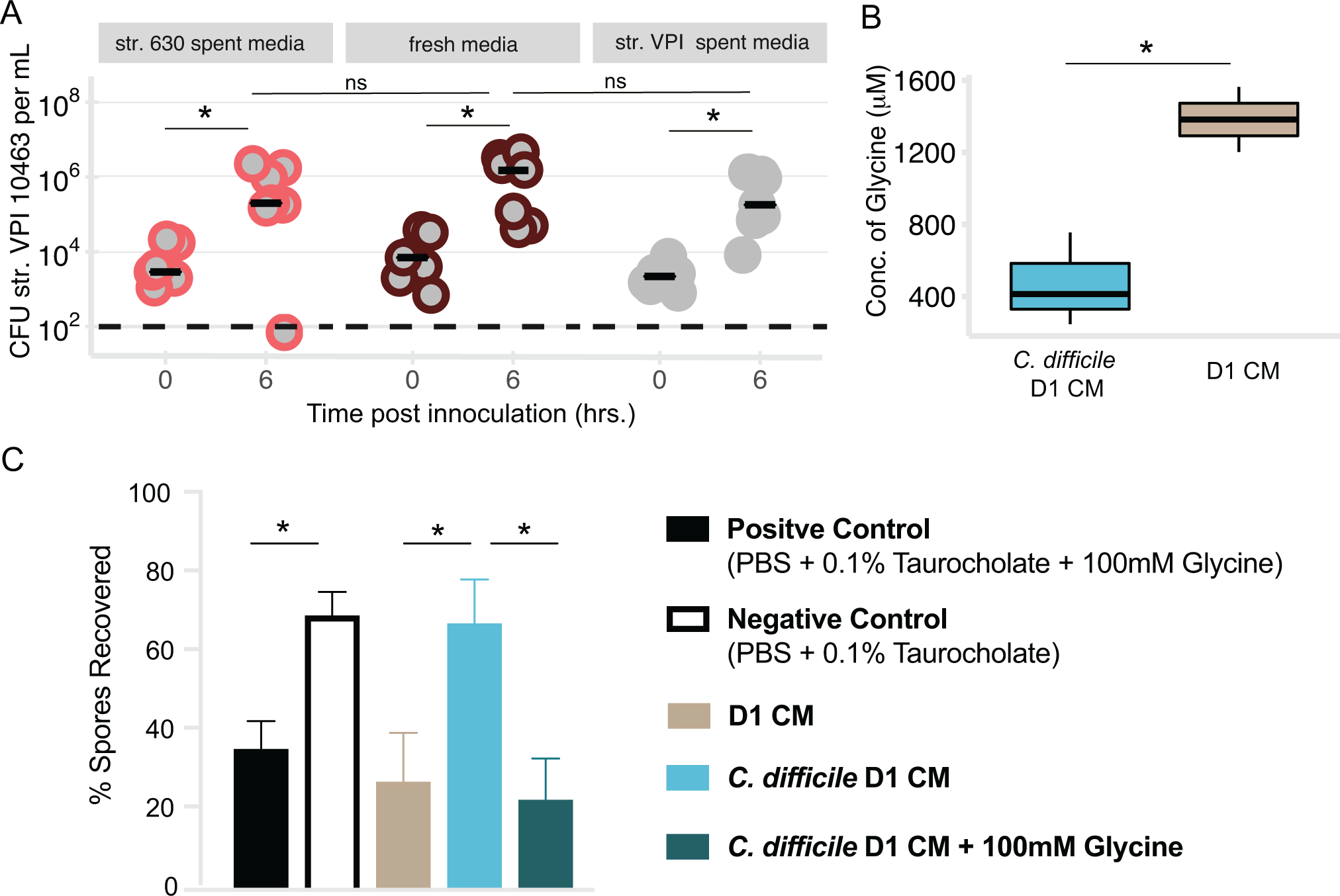
Colonization with *C. difficile* str. 630 significantly reduces the co-germinant glycine in cecal contents, leading to reduced germination of invading strain. **A**. Spent cecal medium from 24hrs growth of str. 630 *ex vivo* supports robust growth of str. VPI 10463. Filter sterilized spent cecal medium was inoculated with vegetative str. VPI 10463 and colonization was monitored by quantitative culture at time zero and six hrs. Cecal medium that grew str. 630 supported robust growth of str. VPI 10463. Str. VPI 10463 colonization at time (t) = 0 vs. t = 6 hrs. in str. 630 spent culture, *P* < 0.05. Str. VPI 10463 colonization at t =0 vs. t= 6 hrs. in fresh medium, *P* < 0.01. Str. VPI 10463 colonization at t =0 vs. t= 6 hrs. in str. VPI 10463 spent culture media, *P* < 0.01. Data represents two independent experiments using separate batches of cecal medium (pooled from at least 6 mice for each batch) run in at least triplicate. Black bars represent median. **B.** Concentration of glycine (μM) in cecal medium made from cecal contents of mice one-day post infection with *C. difficile* (*C. difficile* D1 CM) compared to cecal medium made from mock challenged mice (D1 CM), *P* < 0.05. Data represent values from two separate batches of cecal medium per group. **C.** *Ex vivo* germination of str. VPI 10463 spores in cecal medium from panel B. PBS + 0.1% sodium taurocholate served as a negative control while PBS supplemented with 0.1% sodium taurocholate and 100mM glycine served as a positive control. Bars represent mean with SEM. Negative control vs positive control, *P* < 0.05. D1 CM vs C. *difficile* D1 CM, *P* < 0.05. *C. difficile* D1 CM vs. *C. difficile* D1 CM + 100mM glycine, *P* < 0.05. Data are from at least two independent experiments run in at least duplicate. Statistical significance was calculated using a Wilcoxon test with Benjamini-Hochberg correction (A), t-test (B), or ANOVA with Sidak’s multiple comparisons test (C).

Our group previously observed that 18 hrs. of colonization with strain 630 had significantly decreased levels of amino acids relative to susceptible mice (27). Vegetative *C. difficile* can utilize certain amino acids to enhance growth (28–30). Some of the same amino acids used for growth can also serve as spore co-germinant (31). Since the primary infectious form of *C. difficile* is the not the vegetative cell but rather the environmentally stable spore, we hypothesized that decreased amino acids might mediate diminished levels of germination and thus limit colonization by the invading strain (32). Using targeted metabolomics, we measured concentrations of amino acids in cecal medium (CM) made from mice that had been off of cefoperazone for three days (D1 CM) compared to mice that had been off of cefoperazone for two days followed by infection with *C. difficile* for 24 hrs. (*C. difficile* D1 CM). 24 hrs. of colonization with str. 630 (or a non-toxigenic strain) results in a significant reduction of glycine relative to mock challenged susceptible animals (figure 5B, supplemental figure 6). Glycine is one of the most effective amino acid signals for germination of *C. difficile* spores (33). As we had already ruled out limitation of vegetative growth, we asked if this decrease in glycine altered germination of str. VPI 10463 spores. Germination was assessed by incubation of spores in a given condition for 15 minutes followed by heat-treatment to kill any cells that germinated. If germination occurred, then the post-heat CFU would be lower than the pre-heat amount; if there was minimal germination, the spores would survive heating and levels would remain constant between the pre and post time points.

Heating the spores incubated in PBS+ 0.1% taurocholate + 100mM glycine resulted in a significant reduction in CFU with only 35% of spores recovered, indicating robust germination (figure 5C). However, heating spores incubated PBS + taurocholate resulted in minimal changes between the pre- and post-levels with 70% of spores recovered, suggesting minimal germination. *C. difficile* str. VPI 10463 spores incubated in D1 CM resulted in recovery of 27% of spores, suggesting that this medium supports germination. However, when spores were incubated in *C. difficile* D1 CM, 67% of spores were recovered suggesting that this medium does not support robust germination. *C. difficile* D1 CM enabled significantly less germination compared to D1 CM (*P* < 0.05). To determine if the observed decrease in germination of str. VPI 10463 spores in *C. difficile* D1 CM was due to lower glycine we tested if addition of exogenous glycine could rescue germination. Addition of 100mM glycine to *C. difficile* D1 CM significantly decreased the percent of spores recovered compared to *C. difficile* D1 CM alone (*P* < 0.05).

To provide *in vivo* support for the hypothesis that the protection against infection with VPI 10463 afforded by pre-colonization with str. 630 was due to decreased germination of the more virulent strain, we tested if challenging mice with vegetative str. VPI 10463 overcame this colonization resistance. SPF mice were made susceptible to infection and then challenged with str. 630 spores, the following day the mice were challenged with either 6 x10^4^ CFU of str. VPI spores or 1 x10^7^ CFU of str. VPI 10463 vegetative cells (Supplemental figure 7A). We used a higher dose of vegetative cells because in our hands, over 1 x10^5^ CFU of vegetative cells is necessary to generate a severe infection within two days of challenge, the timeline we were observing with our previous experiments in this study (13). Two days after challenge with the lethal strain, mice given vegetative str. VPI 10463 cells had a modest but significantly higher clinical score compared to animals challenged with spores (Supplemental figure 7B). Together these results demonstrate that pre-colonization with str. 630 reduces levels of glycine in the gut leading to a reduction in the ability of a second strain to germinate.

## Discussion

The role of the gut microbiota in limiting colonization by *C. difficile* has been appreciated for over three decades, however how the microbiota provides colonization resistance remains to be fully elucidated (12, 23). Many studies have focused on a top-down approach to identify and create defined consortia that confer the same protection as the intact community (34, 35). We took an alternative approach and built off the observation that administration of a single bacterium (non-toxigenic *C. difficile*) limited subsequent colonization by another strain of *C. difficile* (10, 36). We sought to determine the mechanisms by which pre-colonization with one strain of *C. difficile* protects from infection with another. We hypothesized that protection was the result of both intraspecific bacterial competition and the development of host immunity to *C. diffic*ile antigens, including the toxins. To evaluate this, we utilized two well-characterized lab strains that, despite being differentially virulent in our mouse model, express nearly identical forms of both TcdA and TcdB (37).

Using multiple infection models we determined that pre-colonization with a less virulent strain is sufficient to protect from challenge with a lethal strain of *C. difficile,* even in the absence of adaptive immunity. Additionally, we showed that protection is dependent on high levels of colonization by the less virulent strain and str. 630 alone is sufficient to limit colonization of the invading strain. While we observed complete exclusion of str. VPI 10463 in the gnotobiotic mice, since we were unable to directly monitor levels of str. VPI 10463 in our SPF mice, we were unable to determine if pre-colonization with str. 630 completely excludes str. VPI 10463 in the context of a more complex microbiota. The prevailing hypothesis has been that pre-colonization with one strain of *C. difficile* limits vegetative growth of the challenging strain. Our results question this model, as 24 hrs. of growth by one strain is not sufficient to deplete nutrients such that it prevented vegetative growth of a second strain.

While other bacterial therapies for *C. difficile* infection such as fecal microbiota transplants can lead to clearance of *C. difficile* from the gut, we were unable to use str. 630 to “treat” mice already colonized with the lethal strain (Supplemental figure 8) (6). Recently, another group reported that a different toxigenic but low virulence strain of *C. difficile* could both protect and “treat” infection with str. VPI 10463 (25). This highlights an important consideration when designing bacterial based therapies for treatment of CDI, as unique strains have different relative fitness. The ability to detect low levels of str. 630 in our co-inoculation model suggests that the strains are able to segregate niches, however more work will be needed to fully elucidate which nutrients are preferred by each strain. Additionally, experiments focused on development of microbial therapies to restore colonization resistance against *C. difficile* should seek to understand the metabolic pathways that enable certain strains to outcompete established *C. difficile* versus inhibit colonization of invading strains as has been done with *E. coli* in the streptomycin mouse model (38, 39).

The major finding from this study is that reduction of amino acids, specifically glycine, following colonization with one strain of *C. difficile* is sufficient to decrease germination of the second strain. This suggests that the axis of intraspecific competition for nutrients includes multiple life-stages of *C. difficile*. Although this type of indirect competition across multiple life stages has been well recognized in macroecology, to our knowledge it has not yet been applied to the study of *C. difficile* infection (40, 41). To date, studies evaluating the role of the microbiota in altering germination of *C. difficile* spores *in vivo* have primarily focused on bile acids, specifically the primary bile acid taurocholate and the secondary bile acid deoxycholate (2, 34, 35). While bile acids play important roles in germination, vegetative growth, and toxin activity, this work demonstrates that microbial metabolism of other nutrients can also affect germination (42). Furthermore, these results suggest that targeting nutrients used by all life stages including those that are metabolically inactive (such as a spore) could be an improved strategy when developing bacterial therapeutics that aim to restore colonization resistance in the gut.

## Materials and Methods

### Animals and housing

Both male and female mice aged five to twelve weeks were used in these studies. The wild-type (WT) C57BL/6 specific-pathogen-free (SPF) mice were from a breeding colony originally derived from the Jackson Laboratory nearly 20 years ago. The RAG1^-/-^ (B6.129S7-*Rag1^tm1Mom^*/J) SPF mice were from a breeding colony started with mice from the Jackson Laboratory in 2013. Germfree RAG1^-/-^ mice were obtained from a colony established and maintained by the University of Michigan germ free facility.

Animal husbandry was performed in an AAALAC-accredited facility. Animals were housed with autoclaved cages, bedding, and water bottles. Mice were fed a standard irradiated chow (LabDiet 5LOD) and had access to food and water *ad libitum*. Cage changes were performed in a biological safety cabinet. To prevent cross-contamination between cages, hydrogen peroxide-based disinfectants in addition to frequent glove changes were utilized during all manipulation of SPF animals. A chlorine-based disinfectant was used during manipulation of the germfree mice. The frequency of cage changes varied depending on the experiment. All mice were maintained under a cycle of 12-hours (hrs.) of light/dark in facilities maintained at a temperature of 72° C +/− 4 degrees. Animal sample size was not determined by a statistical method. Multiple cages of animals for each treatment were used to control for possible differences in the microbiota between cages. Mice were evaluated daily for signs of disease, those determined to be moribund were euthanized by CO_2_ asphyxiation. The University Committee on the Care and Use of Animals at the University of Michigan approved this study.

### Preparation of spore or vegetative inocula of *C. difficile*

Spore stocks of *C. difficile* strains 630 (ATCC BAA-1382) and VPI 10463 (ATCC 43255) were prepared as previously described (15).

Vegetative *C. difficile* was cultured and manipulated in a vinyl anaerobic chamber (Coy Laboratory Products) at 37°C. To generate the vegetative cell inoculum, strain VPI 10463 was streaked from the spore stock on to a plate of pre-reduced cycloserine-cefoxitin-fructose agar containing 0.1% taurocholate (TCCFA). TCCFA was prepared as previously described (15). The following day a colony from the TCCFA was sub-cultured on to brain-heart infusion agar (BHI). After overnight growth, a colony from this plate was inoculated into 5mL of brain-heart infusion broth supplemented with 0.01% cysteine (BHIS) and incubated overnight. The following morning, broth-grown bacteria were harvested by centrifugation at 4000xg for 13 min. The supernatant was discarded, and the pellet was resuspended in 5 mL of sterile phosphate buffered saline (PBS) (Gibco, 10010023). The pellet was washed two more times before being used to challenge mice.

### Preparation of autoclaved *C. difficile*

Heat-killed *C. difficile* strain 630 was made from an overnight culture grown at 37°C in BHIS broth. The culture was enumerated by plating for colony forming units (CFU) per mL^-1^. Broth-grown bacteria were harvested by centrifugation and the cell pellet was washed and resuspended in PBS, at a density of 2.7×10^10^ CFU per mL^-1^. The suspension was autoclaved at 121°C and 14 psi for 30 min to kill the vegetative cells and inactivate any spores. In experiments using heat-killed *C. difficile,* mice received 10^9^ CFU equivalents in 0.05mL. A portion of the sample was cultured on pre-reduced TCCFA to confirm that the inactivation was successful.

### Infections

In experiments using both WT and RAG1^-/-^ SPF mice, age and sex matched mice were co-housed starting at three weeks of age for 33 days through antibiotic administration. Upon infection, animals were separated into single genotype housing.

All SPF mice received the antibiotic cefoperazone (MP Pharmaceuticals, 0219969501) dissolved in Gibco distilled water at concentration of 0.5 mg/mL administered ad libitum for 10 days (15). While mice were on antibiotics, the water was changed every two days. Following completion of antibiotics, mice were given plain Gibco distilled water for two days before challenge with either spores or water (mock). *C. difficile* spores suspended in 50-100μL of Gibco distilled water were administered via oral gavage. The number of viable spores in each inoculums was enumerated by plating for CFU per mL^-1^ on pre-reduced TCCFA. Mice were infected with between 10^3^-10^4^ spores of strain 630. Over the course of the infection, mice were weighed routinely and stool was collected for quantitative culture.

In our long-term colonization model, 41 days after primary infection mice were given an intraperitoneal injection (IP) of clindamycin (Sigma, C5269) in sterile saline at concentration of 10mg/kg to perturb the gut microbial community as described previously (6, 13). The next day mice were either mock challenged with water or with between 10^4^-10^5^ spores from str. VPI 10463. Mice were euthanized two days post infection with str. VPI 10463 (day 44 of the model) and samples for quantification of colonization, toxin cytotoxicity, and histopathology were collected.

In the short-term infection model, mice were challenged with spores, heat-killed str. 630 or vehicle on day 0 and the following day challenged with spores or vegetative cells of strain VPI 10463 (see Supplemental figure 2 and 7D for experiment specific timelines).

In the simultaneous co-infections, mice were challenged with different ratios of str. 630 and str. VPI 10463 within a total amount of 10^4^ spores. During infections when indicated mice were scored for clinical signs of disease based on criteria previously described by (43).

At the conclusion of each experiment, mice used in dual genotype experiments were genotyped using DNA from an ear snip using primers and cycling conditions as outlined by The Jackson Laboratory.

### Quantitative culture from intestinal content

Fecal pellets or colonic content were collected from each mouse into pre-weighted sterile tubes. Following collection, the tubes were reweighed and passed into an anaerobic chamber (Coy Laboratories). In the chamber, each sample was diluted 1 to 10 (w/v) using pre-reduced sterile PBS and serially diluted. 100uL of a given dilution was spread on to pre-reduced TCCFA or when appropriate TCCFA supplemented with either 2 or 6ug/mL of erythromycin (Sigma, E0774). Strain 630 is erythromycin resistant; use of 2ug/mL of erythromycin in TCCFA plates reduced background growth from other bacteria in the sample while TCCFA with 6ug/mL of erythromycin enabled selection of str. 630 (erythromycin resistant) from str. VPI 10463 (erythromycin sensitive). Plates were incubated at 37°C in the anaerobic chamber and colonies were enumerated at 18-24 hrs. Since taurocholate was used in these plates, the colony counts represent both vegetative cells and spores in each sample. Plates that were used to determine if mice were negative for *C. difficile* were held for 48 hrs.

### Toxin cytotoxicity assay

Intestinal content was collected from each mouse into a pre-weighted sterile tube and stored at -80°C. At the start of the assay each sample was diluted 1:10 weight per volume using sterile PBS. Following dilution, the sample was filter-sterilized through a 0.22μm filter and the activity of the toxins was assessed using a Vero cell rounding-based cytotoxicity assay as described previously (6, 44). The cytotoxicity titer was determined for each sample as the last dilution which resulted in at least 80% cell rounding. Toxin titers are reported as the Log_10_ of the reciprocal of the cytotoxicity titer.

### Histopathology evaluation

Mouse ceca and colon tissue were saved in histopathology cassettes and fixed in 10% formalin, followed by storage in 70% ethanol. McClinchey Histology Labs Inc (Stockbridge, MI) prepared the tissue including embedding samples in paraffin, sectioning, and generation of haematoxylin and eosin stained slides. A board-certified veterinary pathologist scored the slides blinded to the experimental groups, using previously described criteria (13, 44).

### Anti-TcdA IgG ELISA

Blood was collected from mice via saphenous vein puncture into capillary blood collection tubes. Samples were spun down and serum was stored at -80°C. Levels of IgG specific to *C. difficile* TcdA were measured in serum from mice using a previously described ELISA protocol (15) with a few modifications. Briefly, 96-well EIA plates were coated with 100μl of purified *C. difficile* TcdA (List Laboratories, 152C) at 1μg/mL in 0.05M sodium bicarbonate buffer pH 9.6 overnight at 4°C. Mouse serum was diluted to 1:400 in blocking buffer and serially diluted 1:3. Each sample was run in duplicate. Negative controls included pre-immune sera from the mice as well as wells serum negative wells. A positive control consisting of a monoclonal mouse anti-TcdA IgG antibody was run on each plate. The optical density at 410 nm was recorded on a VersaMax plate reader (Molecular Devices, Sunnyvale CA). The anti-Toxin A (TcdA) IgG titer was defined as the last dilution where both replicates had an OD_410_ greater than mean absorbance of all negative wells on the plate plus three times the standard deviation from that mean.

### Serum neutralizing anti-toxin antibody assay

The serum neutralizing anti-toxin antibody assay was based on a previously described assay (7). With the following modifications: the concentration of purified TcdA or TcdB (List Laboratories) that resulted in 100% cell rounding was determined empirically, using the toxin activity assay. Vero cells were seeded at a density of 10^5^ cells/0.075mL per well onto tissue culture treated plates. Four times the concentration of toxin that resulted in 100% cells rounding (64μg/mL for TcdA and 4μg/mL for TcdB) was incubated with serial dilutions of serum from mice. The serum toxin mixture was added to Vero cells and incubated overnight. The neutralizing titer was determined to be the last dilution which had less than 50% of round cells in the well. Each sample was run in duplicate and the results from discordant wells were averaged. Toxin only wells served as negative controls while goat anti-TcdA and B serum served as a positive control (TechLab,T5015). Sera from mice before they were infected were run as an additional negative control.

### Quantitative PCR

Primer3 (45) was used to design primers that differentiated strain VPI 10463 (IMG Genome ID: 2512047057) from strain 630 (GenBank: AM180355.1) using publicly available genomes. While the primers differentiated between the strains in pure culture they also picked up other bacteria in the context of the microbiota and were only used in gnotobiotic experiments. *C. difficile* str. VPI 10463 primers: Forward: 5’- TTTCACATGAGCGGACAGGC -3’, Reverse: 5’- TCCGAAGGAGGTTTCCGGTT-3’. The expected product size is 153 nucleotides and the optimal annealing temperature was empirically determined to be 56°C.

For quantitative PCR, DNA from fecal samples were diluted in ultrapure water such that 20 ng of DNA was added to each reaction. Each sample was run in triplicate. Samples were loaded into a Light cycler 480 multiwell plate (Roche, 04729692001), with FastStart Essential DNA Green master mix (Roche, 06402712001), and 0.5μM of each primer. The plate was sealed with optically clear sealing tape, briefly spun down, and run on the Roche LightCycler 96. The following conditions were used for qPCR: 95°C for 10 minutes, followed by 30 cycles of 95°C for 10 seconds, 56°C for 10 seconds, and 72°C for 10 seconds. A melt cycle was run at the conclusion of the amplification.

Genomic DNA from str. VPI 10463 from a culture with a known quantity of CFU was used to generate a standard curve. Negative controls included no template control wells in addition to dilutions of genomic DNA from str. 630.

### Preparation of cecal medium

Cecal medium was prepared from SPF mice that were treated for ten-days with cefoperazone. Uninfected mice were sacrificed two (day 0 post challenge) or three days (equivalent to day 1 post challenge) after the cessation of the antibiotics. Cecal medium was also made from infected mice, following our antibiotic regimen, two days after the completion of antibiotics mice were infected with *C. difficile* and sacrificed one day later (*C. difficile* day 1). Following euthanasia, the cecum was removed and the content was harvested into a sterile 50mL conical. The tube was spun at 3000 rpm at room temperature (RT) for 10 minutes to separate the solid matter from the liquid portion. The liquid portion of the sample was removed and diluted 1:2 by volume in sterile PBS without calcium or magnesium (Gibco, 10010023). The sample was spun again at 3000 rpm for 5 minutes at RT; the liquid portion was then sterilized using filters with successively smaller pore size (from 0.8μm, to 0.45μm, and finally a 0.22μm filter). Following passage through the 0.22μm filter, the medium was frozen at -80°C until use. The medium was tested for sterility by inoculating into pre-reduced BHI in the anaerobic chamber; samples that gave rise to turbid growth after 48 hrs. were discarded.

### *Ex vivo* vegetative growth

To determine if 24 hrs. of str. 630 growth depletes the gut of the nutrients required for vegetative growth of str. VPI 10463 we used an *ex vivo* approach, utilizing sterile cecal medium, prepared as described above, from susceptible mice. 180μL aliquots of day 0 cecal medium were thawed and allowed to equilibrate in the anaerobic chamber overnight. The inocula were prepared from an overnight culture of *C. difficile* (strains 630 and VPI 10463) grown in BHI + 0.01% cysteine. Each culture was back diluted 1:10 and incubated at 37°C for two hrs. After two hrs. had elapsed, 500μL of the culture was spun down for four minutes in the anaerobic chamber using a mini-centrifuge. To prevent the introduction of nutrients from carry over BHI, the supernatant was removed and the tubes were spun for an additional minute. Any remaining BHI was removed and the pellets were then suspended in 500μL of sterile anaerobic PBS. Both strains were then diluted 1:100 into sterile anaerobic PBS. 20μl of this 1:100 dilution was inoculated into the cecal medium. Vegetative CFU were enumerated at T=0, 24 hrs. by plating on cycloserine-cefoxitin-fructose agar (CCFA, which lacks the germinant taurocholate). Additionally, samples were plated on BHI + 0.01% cysteine to check for any contamination. After 24 hrs., the samples were removed from the chamber; cells were pelleted by centrifugation at 2,000 g for five minutes. The supernatant was sterilized by passage through a 0.22μm 96-well filter plate and the sterile flow through was passed back into the chamber to equilibrate overnight. The following day, the VPI 10463 inoculum was prepared as described earlier and inoculated into the spent cecal medium. CFU were monitored by plating samples at T=0 and 6 hrs. on CCFA and BHI+0.01% cysteine.

### *Ex vivo* germination assay

To determine if 24 hrs. of strain 630 growth alters the ability of VPI 10463 spores to germinate we performed an *ex vivo* germination assay based on a previously described method (46) with the following modifications. Rather than intact content, we used sterile cecal medium from mice that were off of antibiotics for three days (D1 cecal medium) or infected with strain 630 for 24 hrs. (*C. difficile* D1 cecal medium). 180μL aliquots of the cecal medium were thawed and allowed to equilibrate in the anaerobic chamber overnight. The next day 10uL of spores from strain VPI 10463 were inoculated into the medium. Controls included sterile PBS (Gibco, 10010023) + 0.1% sodium taurocholate (Sigma, T4009), and PBS + 0.1% sodium taurocholate + 100 mM glycine (Sigma, 50046). Following inoculation, samples were incubated anaerobically at RT for 15 minutes, after which approximately half the volume was immediately plated on BHI + 0.1% sodium taurocholate + 0.01% cysteine agar plates and the tubes were passed out of the chamber and placed in a water filled heat block set at 65°C for 20 minutes. The heating step kills off any vegetative cells. Additional controls included plating the spore inoculum on BHI without taurocholate to check for presence of any vegetative cells in the stock as well as heating suspensions of vegetative cells to confirm efficacy of heat killing. Following the 20-minute incubation, tubes were passed back into the chamber and the remaining sample was plated on BHI + 0.1% sodium taurocholate + 0.01% cysteine agar plates. The CFU recorded from the pre-heat plate represents the entire inoculum including remaining spores and any cells that germinated, while the post-heat plate represents only remaining spores. Percent spores-recovered was calculated as the post-heat counts divided by pre-heat count multiplied by 100.

### Metabolomics

Quantification of amino acid concentrations in the cecal medium by targeted metabolomics was performed by the University of Michigan Medical School Metabolomics Core. Amino acids were measured using the Phenomenex EZfaast kit. Samples were extracted, semi purified, derivatized and measured by EI-GCMS using norvaline as an internal standard for normalization as described previously (47).

### Microbial community analysis

Genomic DNA was extracted from approximately 200-300 μl of fecal sample using the MoBio PowerSoil HTP 96 DNA isolation kit (formerly MoBio, now Qiagen) on the Eppendorf EpMotion 5075 automated pipetting system according to manufacturer’s instructions. The University of Michigan Microbial Systems Laboratory constructed amplicon libraries from extracted DNA as described previously (6). Briefly, the V4 region of the 16S rRNA gene was amplified using barcoded dual index primers as described by Kozich et al. (48). The PCR reaction included the following: 5μl of 4μM stock combined primer set, 0.15μl of Accuprime high-fidelity Taq with 2μl of 10× Accuprime PCR II buffer (Life Technologies, 12346094), 11.85μl of PCR-grade water, and 1μl of template. The PCR cycling conditions were as follows: 95°C for 2 minutes, 30 cycles of 95°C for 20 seconds, 55°C for 15 seconds, and 72°C for 5 minutes, and 10 minutes at 72°C. Following construction, libraries were normalized and pooled using the SequelPrep normalization kit (Life Technologies, A10510-01). The concentration of the pooled libraries was determined using the Kapa Biosystems library quantification kit (KapaBiosystems, KK4854) while amplicon size was determined using the Agilent Bioanalyzer high-sensitivity DNA analysis kit (5067–4626). Amplicon libraries were sequenced on the Illumina MiSeq platform using the MiSeq Reagent 222 kit V2 (MS-102-2003) (500 total cycles) with modifications for the primer set. Illumina’s protocol for library preparation was used for 2 nM libraries, with a final loading concentration of 4pM spiked with 10 % PhiX for diversity. The raw paired-end reads of the sequences for all samples used in this study can be accessed in the Sequence Read Archive under PRJNA388335. Raw sequences were curated using the mother v.1.39.0 software package (49) following the Illumina MiSeq standard operating procedure. Briefly, paired end reads were assembled into contigs and aligned to the V4 region using the SLIVA 16S rRNA sequence database (release v128) (50), any sequences that failed to align were removed; sequences that were flagged as possible chimeras by UCHIME were also removed (51). Sequences were classified with a naïve Bayesian classifier (52) using the Ribosomal Database Project (RDP) and clustered into Operational Taxonomic Units (OTUs) using a 97% similarity cutoff with the Opticlust clustering algorithm (53). The number of sequences in each sample was then rarefied to 9,000 sequences to minimize bias due to uneven sampling. Following curation in mothur, further data analysis and figure generation was carried out in R (v 3.6.3) using standard and loadable packages (54).

### Statistical analysis and generation of figures

The data and code for the analysis associated with this study are available at https://github.com/jlleslie/Intraspecific_Competition. For the purpose of distinguishing between values that were detected at the limit of detection (LOD) versus those that were undetected, all results that were not detected by a given assay were plotted at an arbitrary point below the LOD. However, for statistical analysis, the value of LOD/√2 was imputed for undetected values. Figures and statistical analysis were generated in R (v 3.6.3) using standard and loadable packages or GraphPad (Prism version 8). Adobe Illustrator was used to arrange panels and generate final figures.

## Acknowledgements

This work was supported by grants from the National Institutes for Health including F32DK124048 (JLL), T32GM007863 (PT), T32AI007528 (MRB) and U01AI124244 (PDS, VBY). The authors would also like to acknowledge Judith Opp, April Cockburn, and Harriet Carrington of the University of Michigan Microbial Systems Laboratory for constructing and sequencing the amplicon libraries.

In addition, the authors would also like to thank Zach Powers for technical assistance and Sara Poe from the University of Michigan Germ-free Mouse Core for help running the germ-free infections and. Both cores are funded in part by U2CDK110768.

The funding agencies had no role in study design, data collection, analysis, the decision to publish, or preparation of the manuscript.

J.L.L. and V.B.Y. conceived the study. J.L.L.,M.L.J., K.C.V, A.S., M.R.B, T.J.O, L.U., P.T. & I.L.B performed experiments and/or analyzed the data. P.D.S provided insight and feedback on experimental design. All authors contributed to writing the manuscript and had access to the data.

## Supplemental Figures

**Supplemental** **figure 1: Effect of clindamycin on weight and colonization levels A.** Change in weight from the day mice were given clindamycin to the following day. Mock-infected mice (or naïve animals) are a reference point. There was not a significant difference between the infected or mock-infected mice following administration of clindamycin, *P* > 0.05. **B.** *C. difficile* colonization in infected animals one day prior to administration of clindamycin and one day following. Clindamycin significantly increases levels of *C. difficile* str. 630 in colonized mice, *P* < 0.001. For the data included in each figure, statistical significance was calculated using a Wilcoxon test.

**Supplemental** **figure 2: Short-term infection timelines.** A) Schematic of the short-term colonization in SPF mice related to data in main text figures 4A-C. B) Schematic for experiments in germ-free mice related to main text figures 4D-F. C) Schematic for experiments performing simultaneous challenge with different ratios of strains 630 and VPI 10463 spores related to main text figure 4G and supplemental figure 4.

**Supplemental** **figure 3: Validation of primers for str. VPI 10463.**

**A.** Plot of Cq (or crossing-threshold) vs. dilution of genomic DNA from str. VPI 10463, R^2^ = 0.9929. **B.** The resulting PCR products formed one melt peak.

**Supplemental** **figure 4: Plots from co-inoculation experiments show that high levels of strain 630 correlate with protection from disease.**

**A.** Spearman correlation analysis of % of baseline weight vs. Log_10_ CFU of *C. difficile* str. 630 at 2 days post challenge. **B.** Spearman correlation analysis of total clinical score vs. Log_10_ CFU of *C. difficile* str. 630 at 2 days post challenge. **C.** Spearman correlation analysis of % of baseline weight vs. Log_10_ total *C. difficile* CFU on day 2 post challenge. **D.** Spearman correlation analysis of total clinical score vs. Log_10_ total *C. difficile* CFU on day 2 post challenge.

**Supplemental** **figure 5: *Ex vivo* growth in cecal medium.**

**A.** Sterile cecal filtrate from susceptible mice (day 0) was inoculated with vegetative cells of *C. difficile* str. 630, vehicle, or vegetative cells of *C. difficile* str. VPI 10463. Levels of colonization were monitored by quantitative culture at time 0 and 24 hours. Cecal medium supports robust growth of both strains, *P* < 0.001. Data are from two independent experiments run in at least triplicate using separate batches of cecal medium. **B**. Growth of *C. difficile* str. VPI 10463 in cecal medium made from content from mice infected for 24 hrs. with *C. difficile* str. 630 (630 CM). Strain VPI 10463 colonization at t = 0 vs. t = 24 hrs., *P* < 0.05. Statistical significance for all comparisons was calculated using a Wilcoxon test.

**Supplemental figure 6: Targeted metabolomics measuring amino acids in cecal medium from mock challenged or *C. difficile* colonized mice.**

Concentration (uM) of amino acids in cecal medium (CM). D1 CM is medium made from cecal contents of mice that were made susceptible with antibiotics and then mock challenged. *C. difficile* D1 CM is medium from cecal contents of mice that were colonized with a low virulence strain of *C. difficile* for 24 hrs. Data represent at least two different batches of cecal medium per condition, each batch was made from at least 6 pooled cecal content. Statistical significance for all comparisons was calculated using a t-test.

**Supplemental figure 7. Challenge with vegetative str. VPI 10463 abrogates protection by pre-colonization with str. 630.** A) Schematic for experiment challenging mice with vegetative cells or spores of strain VPI 10463 related to supplemental figure 7. Male and female mice were made susceptible with cefoperazone and then pre-colonized with str. 630. The following day, mice were challenged with either spores (n =6, two cages) or vegetative cells (n=7, two cages) of strain VPI 10463. B) Box plots showing clinical signs of disease (a score which encompasses weight-loss, movement, appearance and diarrhea) was significantly elevated on day 3 of the experiment *P* < 0.05. Statistical significance was calculated using a Wilcoxon test.

**Supplemental figure 8. Treatment with strain 630 does not rescue mice previously infected with strain VPI 10463.** Wild-type mice were challenged with strain VPI 10463 spores and the next day either challenged with strain 630 (n= 5) or mock (n=4). Treatment with str. 630 did not protect against weight-loss and mice had to be euthanized. *P* > 0.05, statistical significance was calculated using a Wilcoxon test.

